# Homocysteine thiolactone from translation proofreading is an endogenous ligand for cell-cell signaling by the receptor SdiA

**DOI:** 10.1101/2024.11.10.622854

**Authors:** Lorenzo Eugenio Leiva, Roberto Zuniga, Gregory J. Phillips, Michael Ibba

## Abstract

Several widespread mechanisms enable cellular communication and coordinate the behavior of bacterial populations. The modules responsible for cellular communication are often found in matching pairs: an enzyme synthesizing a signal molecule and a specific receptor capable of decoding it. A notable exception is *Escherichia coli* SdiA, a homolog of known N-acyl homoserine lactone (AHL) receptors for which no corresponding AHL synthase exists. SdiA is an orphan receptor that enables cross-talk by sensing AHLs from other bacterial species. In this work, we investigated if homocysteine thiolactone (HTL), an AHL-like molecule arising from a proofreading reaction of methionyl tRNA synthetase (MetRS) to correct misactivation of homocysteine, participates in *E. coli* communication through SdiA. A comprehensive analysis of MetRS variants producing different levels of HTL showed more extended lag periods due to lower HTL levels, which were restored by HTL supplementation. SdiA mutants abolished the regulation of the transition from the lag phase to the exponential phase induced by HTL. Growth phenotypes were HTL-specific and were not induced by supplementation of other non-endogenous AHLs previously characterized as SdiA ligands. The gene expression of *gadY* and *rmf* regulated by SdiA was modulated by HTL supplementation. Through differential scanning fluorimetry, we show the SdiA-HTL interaction *in vitro*. Together, our observations describe HTL as an SdiA ligand capable of modulating cell-cell communication in bacteria.

## Introduction

Bacteria need to adapt to different environments and stress conditions, and quorum sensing (QS) is a generalized cell-cell bacterial communication mechanism that allows these changes to be synchronized whenever a specific critical density of a diffusible signal molecule is reached (1). The variety of known QS signal molecules has grown over the years. In Gram-negative bacteria, autoinducers type I (AI-I) based on acyl homoserine lactones (AHLs) are the most commonly known class of QS signal molecules (2). They contain an N-acylated homoserine-lactone ring core and a 4-18 acyl chain (3). The features of the aliphatic chains, such as length and some modifications that may be present, give a species-specific character to AI-Is and affect their stability, diffusion, and detection. For most of the bacteria able to produce AI-I, their synthesis is often catalyzed by LuxI-type synthases, which produce AHLs by deriving lactones from S-adenosyl methionine (SAM) and the acyl chain from intermediates of fatty acid metabolism (3).

The most common receptors for QS signals in Gram-negative bacteria are LuxR-type proteins, which are cytoplasmic transcription factors. The formation of the LuxR-autoinducer complex promotes the stability of the receptor and its homodimerization and facilitates subsequent binding to *lux boxes* in the promoter region of target genes (4, 5). Despite the coevolution of the AHL synthase-receptor pair, based on amino acid variation in the binding pocket to give AHL binding specificity, close to 75% of the annotated LuxR homologs correspond to orphan transcriptional factors that are not accompanied by a LuxI synthase counterpart. These receptors are called LuxR-solo class proteins (6). One alternative explanation is that some of these LuxR-solo classes have a relaxed ligand binding specificity that allows recognition of many types of AI-I. For example, QscR is a well-studied LuxR-only receptor from *Pseudomonas aeruginosa*, whose activity can be modulated by C8-HSL, C10-HSL, 3-oxo-C10-HSL, C12-HSL, 3-oxo-C12-HSL and C14-HSL (7, 7a). Furthermore, there are examples of QS related LuxR homologs that respond to non-AHL endogenously produced ligands, for example PluR and PauR which respond to photopyrones and dialkylresorcinol, respectively (7b, 7c). SdiA is an *Escherichia coli* homolog of the LuxR receptor linked to bacterial virulence and tolerance, for which no signaling molecule produced by the same bacteria has been described (8–10). Instead, SdiA responds to ligands produced by other bacteria, a process termed “eavesdropping” (10a, 10b).

Protein synthesis depends on the aminoacyl tRNA synthetases, an essential family of enzymes responsible for defining the genetic code while translating the nucleotide alphabet into proteinogenic amino acids. The aminoacylation process consists of a two-step reaction, where the first involves the recognition and activation of the amino acid before its subsequent transfer to the cognate tRNA. To preserve the fidelity of the process, different mechanisms allow editing of recognition errors resulting from physicochemical similarities between amino acids, such as phenylalanine and tyrosine, branched amino acids, etc (11). These errors can also involve non-proteogenic amino acids, such as D-amino acid enantiomers or intermediates in amino acid biosynthesis. An example of the latter is homocysteine (Hcy), a sulfonated amino acid intermediate in the biosynthesis of methionine (Met) from oxaloacetate in the TCA cycle (12). Although Hcy lacks the final methylation in the R group, methionyl tRNA synthetase (MetRS) can recognize and activate this amino acid. However, the hydrophobic environment of the amino acid binding pocket promotes a highly efficient cyclization reaction between the reactive sulphydryl group of the Hcy and the carbon of the carboxyl group, releasing AMP and producing homocysteine thiolactone (HTL) (13). HTL is a stable molecule at acidic and neutral pH values, is highly diffusible at physiological pH, and has significant similarity to the lactone ring of AHLs (14, 15).

HTL has been considered a so-called carbon leak point for many years because no biological reaction has been described in bacteria to reintegrate HTL into bacterial metabolism. Deleterious effects of HTL accumulation have been described due to the reactivity of the sulfur group and its ability to modify exposed lysines in eukaryotic proteins but not bacteria (16). In this work, we describe how *E. coli* produces HTL mainly via a proofreading reaction of MetRS and how its subsequent accumulation induces cell division without affecting the duplication time. This suggests HTL has the potential to act as a signaling molecule that mimics AHLs. Additionally, we show that this phenotype is dependent on the presence of the LuxR-solo receptor SdiA. The implications of the accumulation and detection of HTL in the bacterial population to regulate the lag period of growth and its implication in bacterial antibiotic tolerance phenomena are also discussed.

## Results

### Changes in the lag period in response to HTL levels

*E. coli metG* is an essential gene since it encodes MetRS, an enzyme responsible for the aminoacylation of both tRNA^fmet^ (tRNA_i_) for translation initiation and tRNA^met^ for elongation. To initiate our studies, we selected a set of *metG* mutants that our previous research showed were deficient in MetRS proofreading activity (17). These strains contain mutations that alter amino acids near the catalytic site of MetRS (*metG83*), delete four amino acids near the anticodon-binding domain (*metETIT*), and truncate the last 47 amino acids that form part of the tRNA binding domain, recapitulating a mutant first isolated in *S. Typhimurium* (*metG630*). When analyzing cell growth, all *metG* mutants showed the same growth patterns as their parental strain in LB medium (Figure 1A). Because LB contains about ∼5.9 mM Met (18), any defect in MetRS proofreading activity could be masked by an excess of the cognate amino acid. Repeating the same analysis in a medium without Met, the strains showed more significant differences in the lag period, but not the doubling time, during the exponential phase of growth (Figure 1B and C).

**Figure 1.**
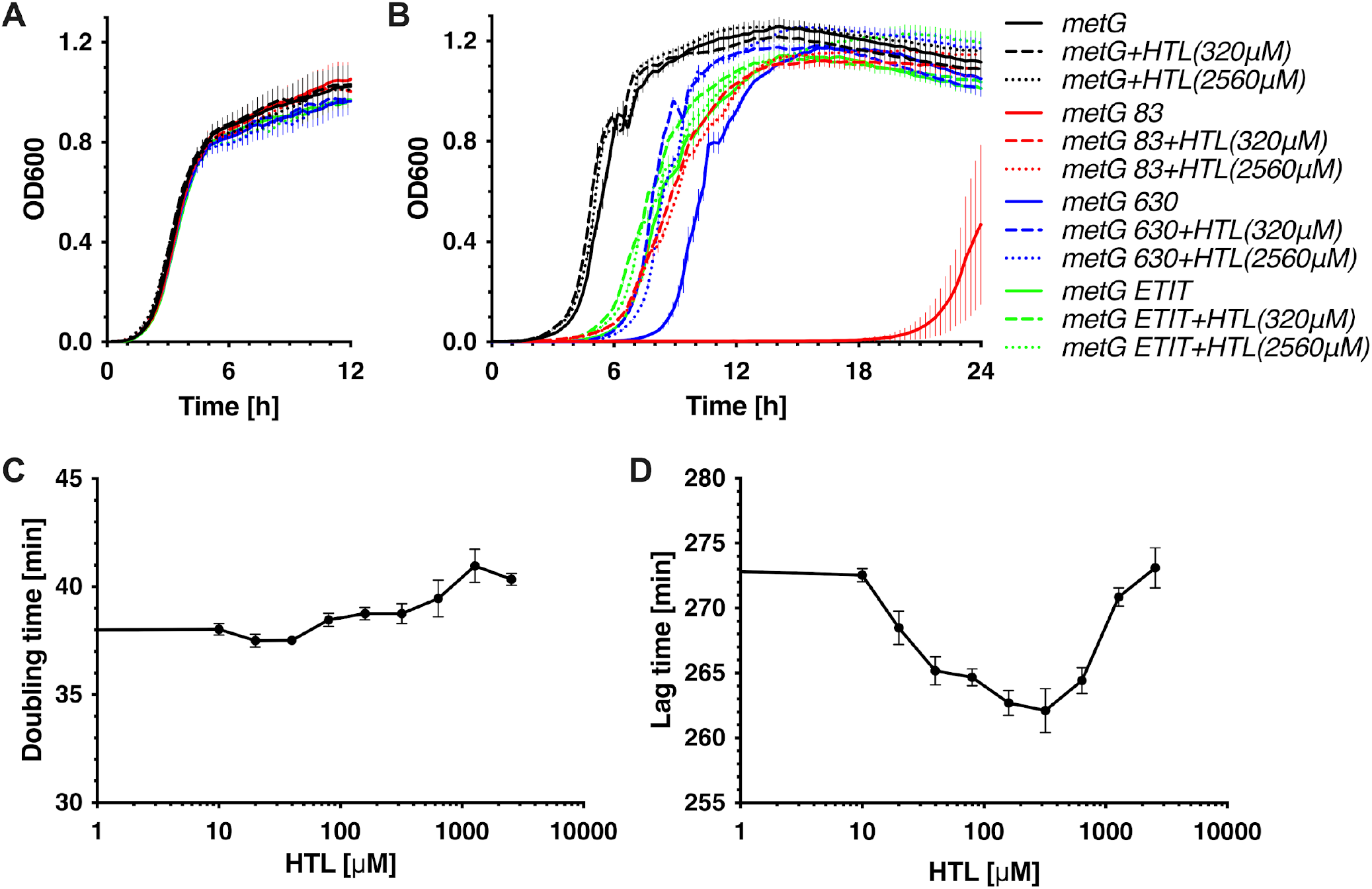
*metG* mutations impact the lag period of growth curves. All strains were grown in LB (A) or M9 medium (B). M9 medium was supplemented with 40μg/ml of each amino acid except Met. The OD at 600nm (OD600) of *E. coli* strains *metG* (wild-type), *metG83, metG630*, and *metG ETIT* was monitored in a microplate reader at 37 °C with constant shaking. Error bars represent standard deviations of biological triplicates. Strains were supplemented with 320 μM or 2560 μM HTL when indicated. The doubling time (C) and lag time (D) were determined from *metG* strain grown with increasing concentrations of HTL.

MetRS, like the isoleucine, leucine, and valine aminoacyl tRNA synthetases, can bind and activate Hcy. However, due to the proofreading activity possessed by these enzymes, Hcy adenylate is rapidly removed from the amino acid binding pocket in a cyclization reaction that releases HTL. To explore whether changes in HTL production rate are related to the differences observed in the lag period, medium without Met was supplemented with HTL. A notable reduction in the lag period was observed in the *metG83* and *metG630 strains*, while a slight effect was observed in the wild type (WT) and *metGETIT* strains (Figure 1B). The reduction in the lag period of the WT gradually decreased to a minimum upon the addition of 320 μM HTL, after which the lag period began to increase (Figure 1D), probably due to the deleterious effects that have been attributed to elevated levels of HTL (19, 20). Higher levels of HTL in *metG83* and *metG630* did not lead to a significant increase in the lag period, suggesting that after a threshold concentration, there is no dose-response effect in decreasing the lag period (Figure 1B). Furthermore, no significant differences in protein synthesis levels were observed between *metG* WT and the mutants (Figure S1). From these results, we can infer that the effects of mutations in *metG* are related to the ability of MetRS to produce and accumulate a critical concentration of HTL.

### Long lag periods correlate with low HTL concentrations

To determine whether strains *metG83* and *metG630* respond to exogenous HTL supplementation due to a deficiency in its synthesis, HTL levels were measured over time. HTL monitoring was based upon two considerations: 1) Because HTL can quickly diffuse through the membrane and no differences in intracellular levels have been observed in Met biosynthesis mutants that produce higher levels of Hcy and HTL, their levels were monitored from the supernatant (16, 21). 2) Because HTL synthesis can be reduced due to amino acid supplementation (22), we used M9 minimal medium where no amino acids were added in cultures or preinocula.

Although this strongly impacted the extension of the growth curves, the same differences remained in the lag periods between WT, *metG83*, and *metG630* strains (Figure 2A). To determine the levels of HTL in the supernatant, we established a fluorometric plate assay based on the reaction between HTL and *ortho*-phtalaldehyde (OPA) (21, 23). We found different dynamics of HTL levels between the strains, where the most rapid accumulation of HTL in the supernatant was observed in the WT, followed by *metG630* and *metG83* (Figure 2B). This is consistent with more extended lag periods in the growth curve being correlated with lower levels of HTL being synthesized by the cell.

**Figure 2.**
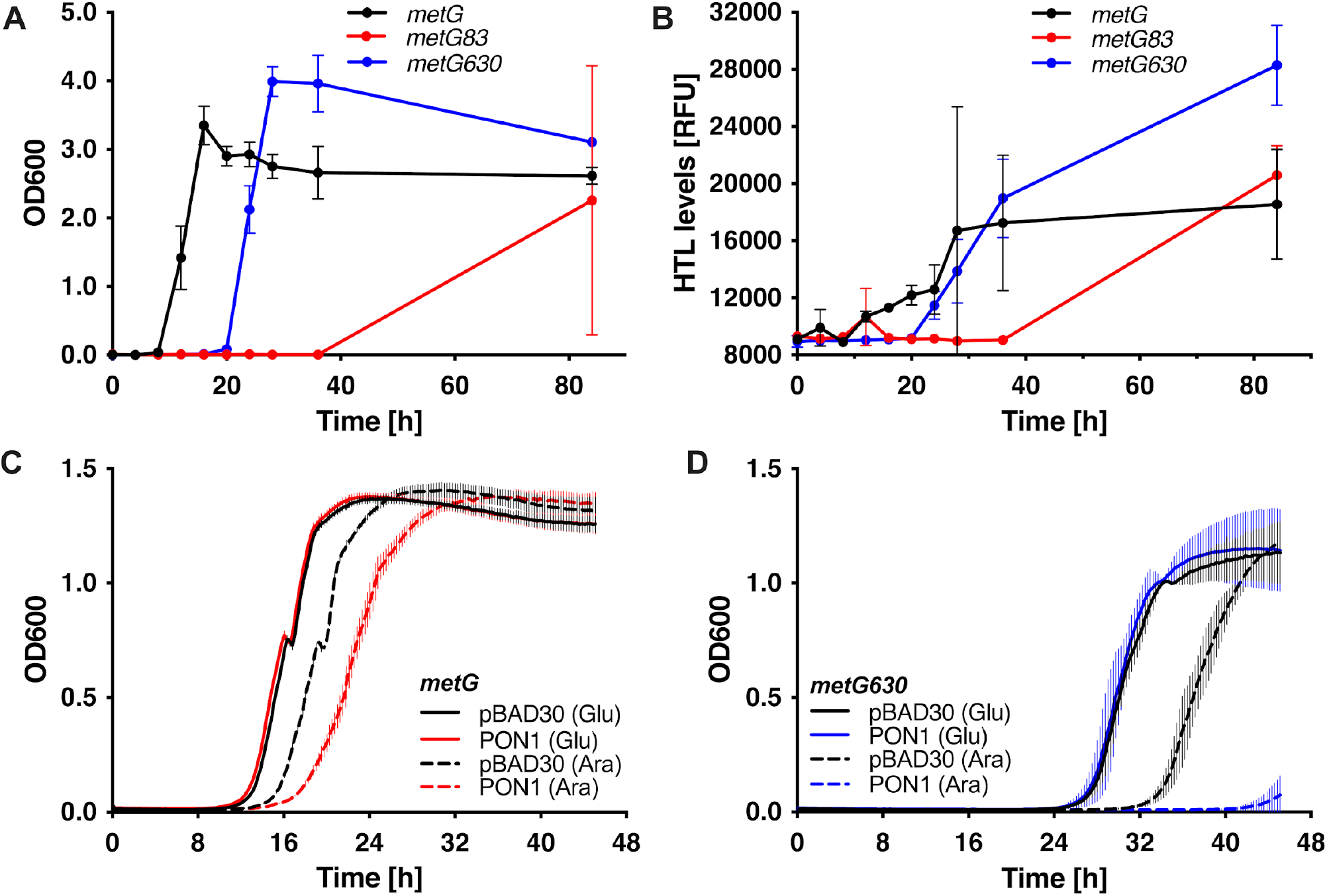
Lag period in response to HTL levels. A) *metG* (wild-type), *metG83*, and *metG630* growth curves in M9 medium without amino acid supplementation (pre-inocula and cultures). Cultures were carried out in test tubes at 37°C with constant shaking. B) Aliquots of the culture supernatants in Figure A were mixed with an orthophotoaldehyde (OPA) solution to produce a fluorogenic derivative of HTL and measure its levels on a TECAN plate fluorimeter. Error bars represent the standard deviation between biological replicates. PON1, a hydrolase that catalyzes the degradation of a wide range of lactones, was cloned into plasmid pBAD30 and transformed into *metG* (wild-type) (C) and *metG630* (D). Strains were grown in M9 medium supplemented with glucose (Glu) or arabinose (Ara) to repress or induce PON1 expression, respectively.

To further corroborate that the effect on culture growth is not an artifact of mutating *metG*, we attempted to reduce HTL levels independently of MetRS proofreading activity. To achieve this goal, we induced the expression of a variant of paraoxonase 1 (PON1), a calcium-dependent hydrolase that catalyzes the degradation of a wide range of lactones. Because the origin of this enzyme is mammalian and shows low solubility in *E. coli*, we used a recombinant variant evolved *in vitro* for bacterial expression (24, 25). The toxic effect of overproduction of PON1 was minimized by cloning it into a low-copy number plasmid, inducible by arabinose supplementation. To investigate the length of the lag period in response to PON1 induction, we compared WT and *metG630* strains transformed with the parental plasmid pBAD30 or the pPON1 plasmid (Figure 2C and D). Both plasmids induced the same growth curve dynamics in WT and *metG630* when the medium was supplemented with glucose to repress expression. However, when cultures were supplemented with arabinose, pPON1-transformed strains showed a more extended lag period before exponential growth, supporting the hypothesis that HTL levels regulate the onset of exponential growth (Figure 2C and D). The more efficient growth observed in the presence of glucose can be explained by its immediate use in glycolysis versus arabinose, which enters metabolism through the pentose pathway.

### Response to HTL and pH in the medium

The finding that HTL can regulate the duration of the lag period raises the question of whether intracellular concentrations of HTL are sufficient to trigger cell division or whether HTL levels coordinate cell division based on its extracellular concentration. Before exploring this last alternative, we must consider the chemistry of the molecule and the changes in the medium during bacterial growth. HTL is a small molecule with a pKa 6.67, which causes it to be primarily uncharged at neutral pH, allowing it to diffuse across membranes (26, 27). HTL has been reported to be very stable under acidic conditions over various temperatures (28). On the contrary, alkaline conditions can induce its degradation by opening the thiolactone ring and reacting with other HTL molecules, forming 2,5-diketopiperazine Hcy (29). This becomes relevant when the pH of the media can show fluctuations depending on the carbon source and oxygenation levels, which can lead to alkalinization of the medium due to amino acid metabolism or acidification due to sugar fermentation (30, 31). Next, to explore whether fluctuation in extracellular HTL levels is more relevant than intracellular HTL levels to trigger cell division, we analyzed the response of HTL supplementation at different pHs in the WT strain. When we looked at the growth curves at pH 7.4, where 84.3% (269.8 μM) of the total supplemented HTL was in neutral form, the lag period was not reduced by HTL supplementation (Figure 3A and B). However, at a pH of 4.2, where only 0.34% (1.1 μM) of the supplemented HTL was neutral, HTL supplementation was necessary to cause a similar lag period to the WT at a pH of 7.4 (Figure 3A and B). Together, these observations suggest that under nutrient-limited growth conditions, the intracellular levels of HTL produced as by-products of Met metabolism are insufficient to trigger exponential growth.

**Figure 3.**
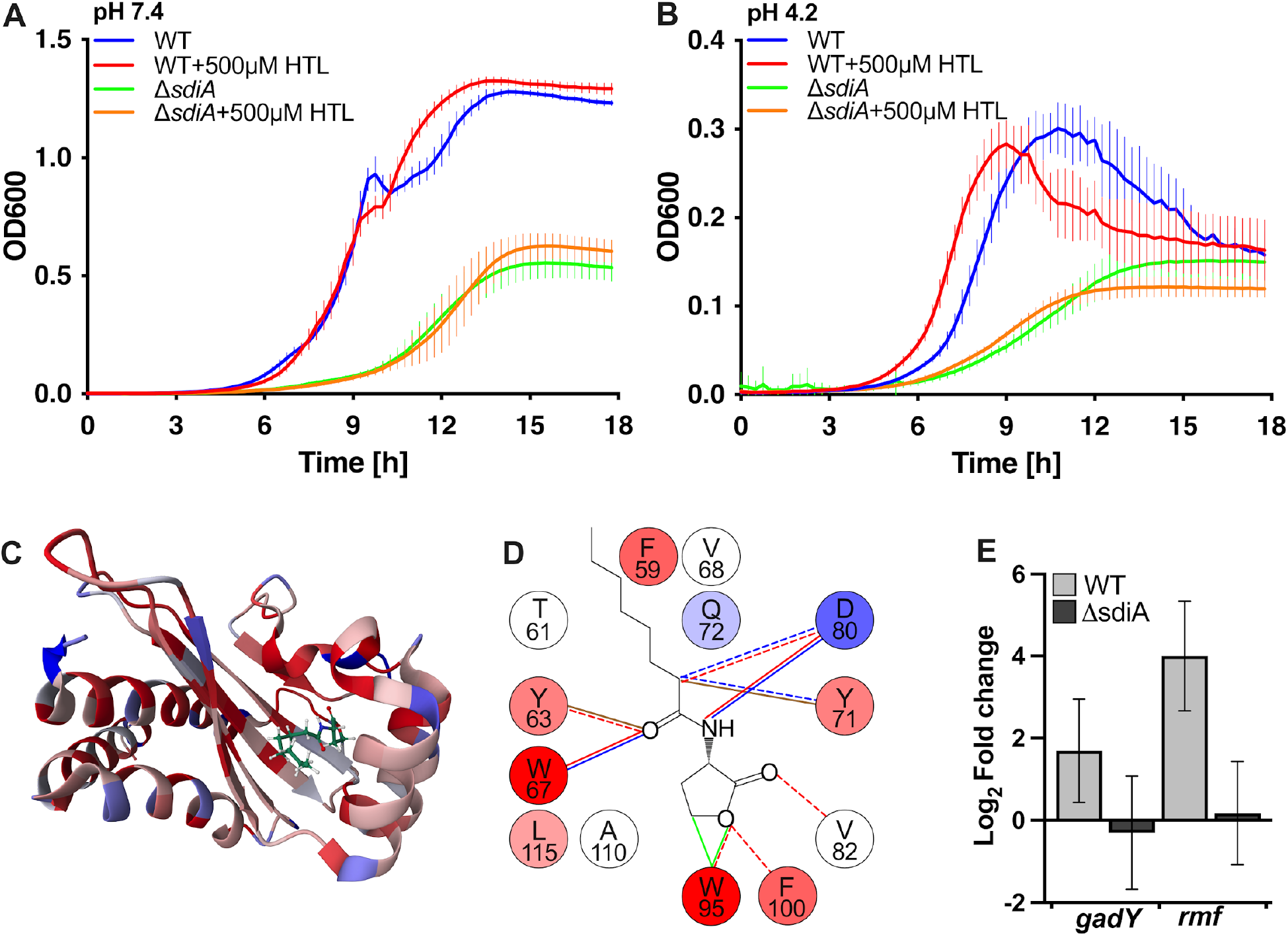
Determination of the role of SdiA in HTL response. *metG* (wild-type) and *metGΔsdiA* (ΔsdiA) were cultured in M9 medium at pH 7.4 (A) and pH 4.2 and supplemented with 500 μM HTL when indicated. The OD at 600 nm (OD600) was monitored in a microplate reader at 37 °C with constant shaking. Error bars represent standard deviations of biological triplicates. (C) The protein structure of a complex between the ligand binding domain of SdiA and the autoinducer N-octanoyl-l-homoserine lactone (C8-HSL) was calculated using NMR data. The colors represent the Hydrophobicity scale: hydrophobic (red) and hydrophilic (blue). D) Description of the autoinducer binding pocket and their interactions: Hydrogen bond (blue lines), polar (red line), Van Der Waals (green lines), hydrophobic (brown lines). Weak interactions are presented with a dashed line. E) Fold change of *gadY* and *rmf* transcripts between cultures in M9 and M9 supplemented with HTL 320 μM for 30 min, for WT and ΔsdiA strains.

### SdiA is required to detect HTL

Although our results showed that HTL is a molecule involved in bacterial signaling, to support that it is part of a communication mechanism, it is necessary to elucidate how HTL is detected. HTL shares a similar structure with the homoserine-lactone ring of AI-I, raising the possibility of crosstalk with QS receptors. SdiA, a homolog of the AI-I receptor LuxR, is present in *E. coli* and has been associated with cell division, antibiotic resistance, virulence in pathogenic strains, biofilm formation, and acid resistance, among other phenotypes (8, 10). While no ligand for SdiA synthesized by *E. coli* has been described to date, its role in detecting AHLs derived from other bacterial species is well documented (e.g. 10a). Consequently, SdiA is a QS receptor that modulates *E. coli* physiology following population changes of different species in the environment (e.g. 10b). Structural studies have analyzed the interaction between SdiA and N-octanoyl-L-homoserine lactone (C8-HSL), showing that many of the most conserved amino acids of the binding pocket interact with the ketone, the amino, and the β carbon of the lactone, all of which are also present in HTL (Figure 3C and D) (5).

To study the role played by SdiA in regulating the lag period dependent on HTL levels, we analyzed the response to HTL supplementation in a Δ*sdiA* mutant *(ΔsdiA::KanR*). Because this *sdiA* mutation itself is sufficient to cause a defect in the growth of *E. coli*, to allow analysis in microplates in a time window no greater than 24h, the effects of neither Δ*sdiA*/*metG83* nor Δ*sdiA*/*metG630* double mutants were investigated. While WT and Δ*sdiA E. coli* showed no response to HTL supplementation at pH 7.4 at acidic pH, where the modulation of the lag period is better observed in WT, the response to HTL addition disappeared in the Δ*sdiA* mutant (Figures 3A and B). This result supports the idea that HTL is a ligand for SdiA.

Several reports that have analyzed the function of SdiA have described targets of this transcription factor, such as *gadY*, which is involved in acidic tolerance, and *rmf*, which encodes a ribosomal modulation factor that coordinates the transition to the stationary phase (8, 32). Because we expect the function of the SdiA transcription factor to be affected by HTL levels, we compared the expression levels of *gadY* and *rfm* from cultures incubated for 30 min in M9 medium with or without HTL (Figure 3E). These results showed an increase in the expression of both genes of ∼4 and ∼20 fold compared to the culture without HTL supplementation. This effect of HTL on *gadY* and *rfm* expression is absent in the Δ*sdiA* mutant, showing that HTL-induced expression changes depend on SdiA (Figure 3E).

To support our model that the observed response to HTL is a consequence of direct interaction with SdiA, we assessed their interaction by using differential scanning fluorimetry to monitor changes in the thermal stability of SdiA in the presence of the ligand (33, 34). Protein stability depends on physical parameters (temperature, salinity, pH, detergents, etc.) and biophysical factors, such as protein-ligand interactions (33, 35). We analyzed the unfolding of SdiA by measuring the fluorescence intensity of SYPRO Orange, which depends on the interaction of the dye with the hydrophobic regions of SdiA exposed during thermal denaturation (Figure 4A). The presence of HTL induced a shift in the unfolding profile to higher temperatures (Figure 4A), increased the apparent melting temperature (Tm_a_), and the temperature that maximized fluorescence for the unfolded protein (Max) (Figure 4 B and C). This change in the Tm suggests that the interaction with HTL stabilizes SdiA.

**Figure 4.**
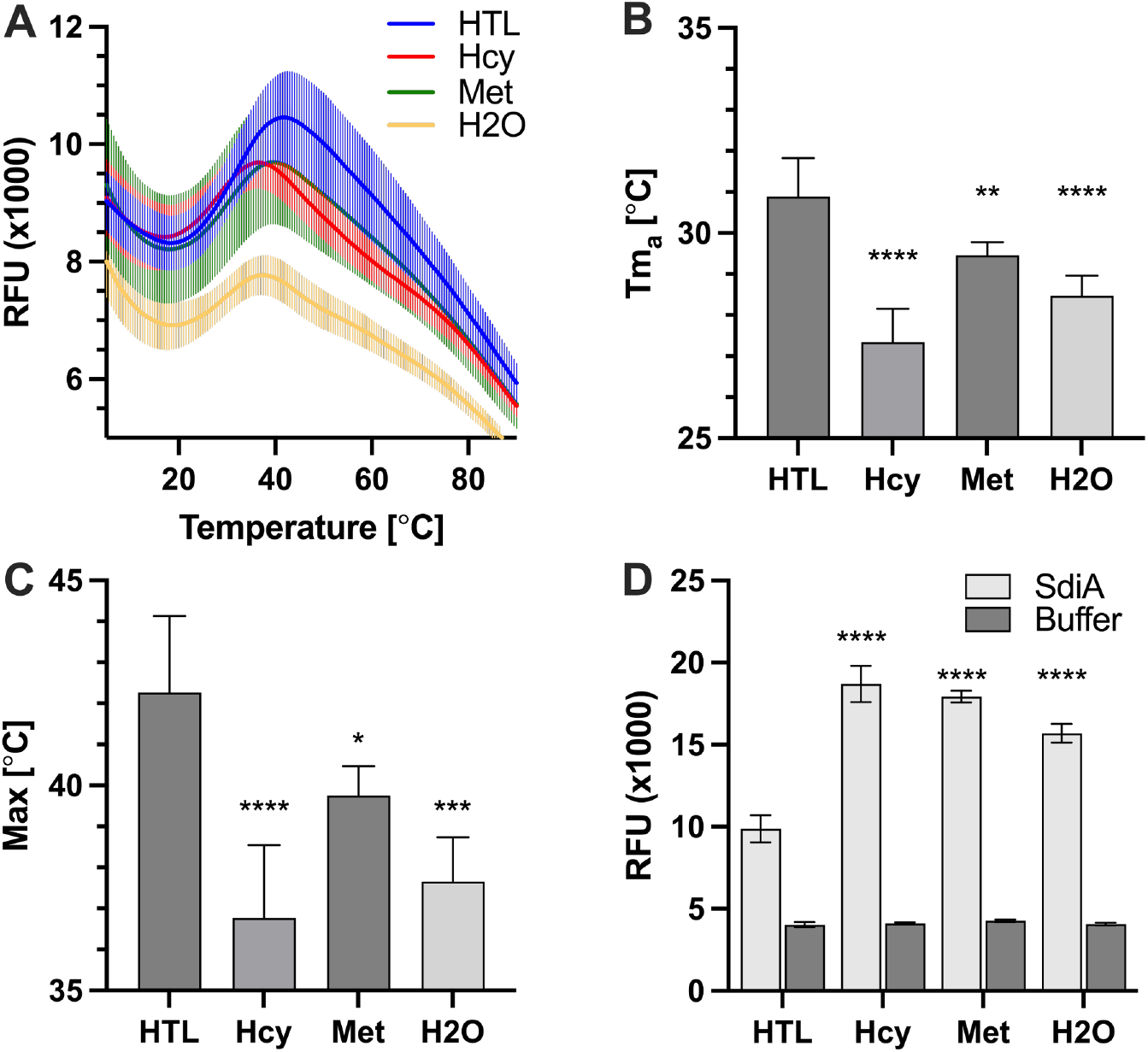
HTL stabilizes SdiA and quenches fluorophore binding to hydrophobic regions. Measurements of SdiA stability in response to HTL, Hcy, and Met were performed using differential scanning fluorimetry. A) SdiA unfolding curve in the presence of SYPRO Orange in a 1:2 ratio. Potential ligands were used at equimolar concentrations (5.4mM). The apparent melting temperature (Tma) (B) and fluorescence maximum (Max) (C) were determined from Figure A. D) Measurement of fluorophore quenching (at a ratio of 1:1.7) at 4°C. The background without protein (buffer) is presented in gray. ****p ≤ 0.0001, ***p ≤ 0.001, **p ≤ 0.01, *p < 0.05, one-way ANOVA with Dunnett versus HTL.

This stabilization effect on SdiA was not observed with Hcy, the cyclization reaction’s precursor amino acid substrate, which shares properties similar to HTL (Figure 4 A, B, and C). Because the effect of HTL supplementation in a Met-rich medium was not observed (Figure 1A), the interaction of SdiA with Met was also evaluated. Although there is a slight increase in melting temperature with the addition of Met, it is less than that observed for HTL. Considering the abundance of hydrophobic amino acids in the ligand binding domain (Figure 3C), the temperature change with Met could result from its greater hydrophobicity. The binding pocket of SdiA provides a highly hydrophobic environment to host the ligand, which also tends to be occupied by the fluorophore during the assay due to its affinity for these hydrophobic regions. When we measured fluorescence immediately after adding the dye to the protein-ligand complex, we observed that only HTL could quench the fluorophore, suggesting that SYPRO Orange rapidly displaces Hcy and Met. At the same time, HTL remains in the binding pocket due to its higher affinity (Figure 4D). Together, these results strongly suggest that HTL plays a regulatory role with SdiA as its receptor.

### Specificity of response to HTL

Prior to this work, little or no activity for SdiA had been reported in the absence of exogenous ligands (35a, 35b). To analyze the selectivity of SdiA for HTL, we compared the effect on the lag period during *metG83* growth following supplementation with equimolar concentrations of a set of HTL analogs (Figure 5). With Met supplementation, we observed a drastic reduction in the lag period, consistent with the rapid growth of the *metG83* mutant previously observed in LB medium (Figures 1 and 5).

**Figure 5.**
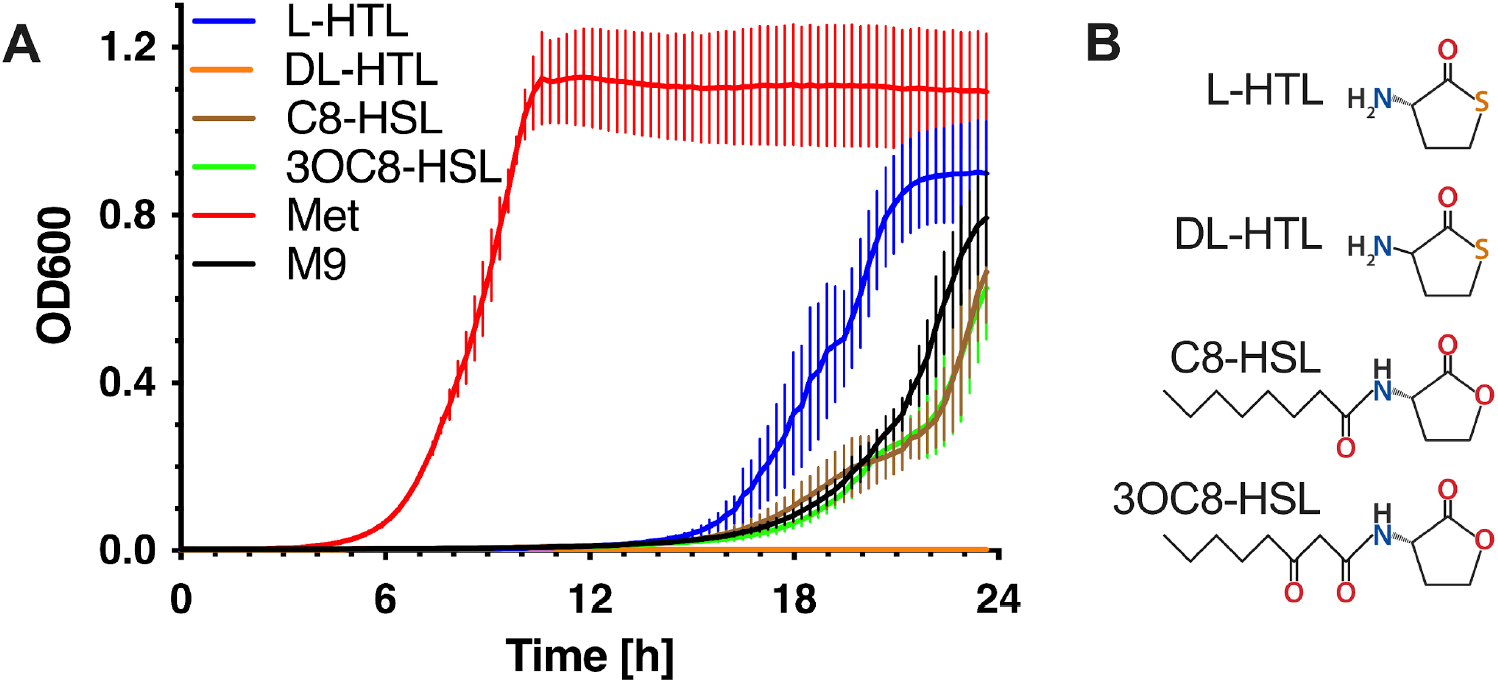
Effect of HTL analogs on the regulation of lag period. A) *metG83* was cultured in M9 medium and supplemented with 320 μM of L-homocysteine thiolactone (L-HTL), methionine (Met), N-octanoyl-L-homoserine lactone (C8-HSL), N-(3-Oxooctanoyl)-L-homoserine lactone (3OC8-HSL) and the racemic mixture of D-HTL and L-HTL (DL-HTL), when indicated. The OD at 600 nm (OD600) was monitored in a microplate reader at 37 °C with constant shaking. Error bars represent standard deviations of biological triplicates. B) HTL homologs added to the culture.

C8-HSL and 3OC8-HSL are autoinducers that have been described in *Burkholderia spp*., *Pseudomonas ssp*., and *Agrobacterium spp*. and were used previously as ligands to study SdiA from *E. coli* (5, 36). We observed that the lag periods following C8-HSL and 3OC8-HSL supplementation are more extended than those for bacteria supplemented with HTL and closer to those grown in M9 medium. The source of HTL in our model is L-Hcy, which will produce L-HTL during MetRS editing activity. However, the DL-HTL mixture was described as an anti-quorum sensing molecule synthesized by coral symbiotic bacteria (37). We found that DL-HTL is insufficient to obtain the critical concentration of L-HTL to reduce the lag period and inhibit growth, suggesting that D-HTL is the enantiomer responsible for the previously reported inhibitory effects of HTL.

## Discussion

### Homocysteine thiolactone is a determinant of lag-to-log phase transition during growth

Here, we describe a previously unrecognized endogenous ligand for SdiA, a LuxR-solo class protein, which senses HTL levels to coordinate *E. coli* growth. Bacteria such as *E. coli* encode the complete Met biosynthesis pathway, which involves the synthesis of Hcy as a precursor. Due to the chemical similarities of Hcy with Met, MetRS recognizes and activates it, forming Hcy adenylate. The proofreading activity of MetRS then catalyzes the cyclization of Hcy to avoid translation errors, generating HTL as a byproduct (Figure 6). Although the accumulation of this molecule was consistently associated with protein homocysteinylation damage (38), we show that HTL accumulation is necessary to modulate the transition between the lag and exponential growth phases (Figure 1 B). This new finding depended mainly on using *metG* mutants, initially identified in experiments designed to select strains resistant or highly tolerant to antibiotics or amino acid analogues. Regulation of the lag period, particularly its extension, is one of the mechanisms described that can induce tolerance to stress conditions and has been shown to be important for antibiotic tolerance (39–41). Given this, the lengthening of the lag period for the *metG* mutants provides a plausible explanation for the high antibiotic tolerance observed for the strains (17).

**Figure 6.**
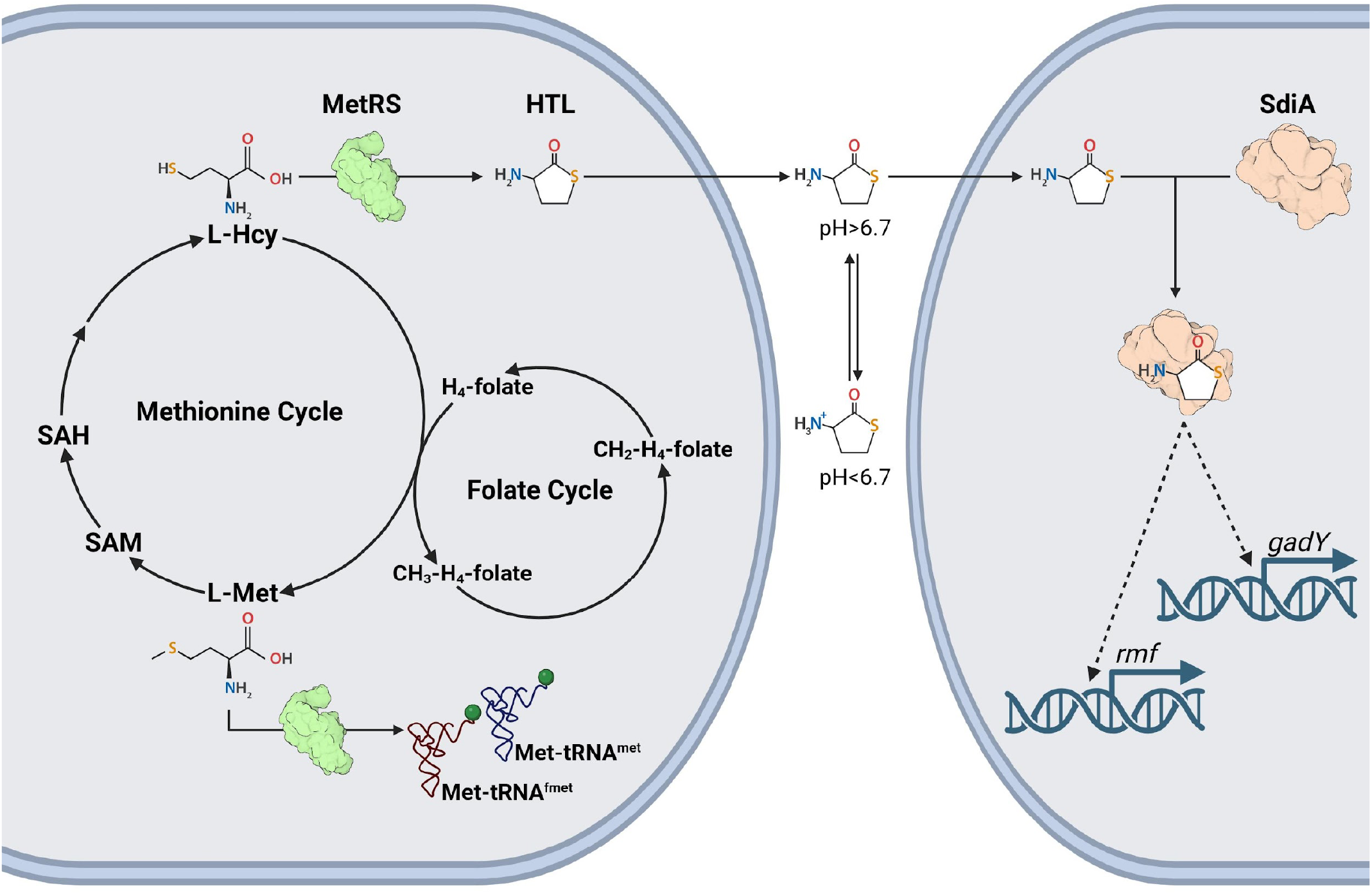
HTL synthesis in response to the balance between L-Hcy and L-Met. The conversion of L-Hcy to L-Met will depend on H4-folate levels. This new L-Met will be used for both translation and synthesis of SAM. Consequently, the balance of L-Hcy and L-Met will depend on the state of the folate cycle, the level of protein synthesis, and the regeneration of L-Hcy in the methionine cycle. Any change in these central processes can induce the accumulation of L-Hcy, its subsequent misactivation by MetRS, and the synthesis of HTL. HTL can quickly diffuse at physiological pH owing to its size and neutrality. After reaching a critical concentration, which will depend on the extracellular pH, HTL enters the cells, activates SdiA, and induces the expression of genes, such as *gadY* and *rmf*.

The extended lag phases observed in the *metG* mutant strains could be primarily correlated to reduced HTL levels rather than a general reduction in protein synthesis rates, as no significant changes in translation activity were observed compared to the parental strain. Also, in an amino acid-limited minimal medium, HTL supplementation reduced the lag time of the growth curves without significantly modifying the doubling time (Figure 1). The central role of HTL was further corroborated in a WT strain when the expression of PON1, an enzyme that hydrolyzes the thiolactone ring forming Hcy, prevented the accumulation of HTL. The same effect was obtained when the permeability of HTL was reduced by using a lower pH in the culture, where the signal molecule gained charge, avoiding the detection of HTL accumulation in the extracellular medium (Figure 3B).

### Homocysteine thiolactone as a signal of amino acid homeostasis

Previous reports have shown how the level of HTL is affected by fluctuations in the balance of intracellular Hcy and Met concentrations. There, higher levels of HTL were detected instead of the higher levels of Hcy that might be expected in mutants of *metE*, one of the two genes involved in the final methylation of Hcy in Met synthesis (22). This suggests that MetRS proofreading prevents the accumulation of excess Hcy that might otherwise disrupt branched-chain amino acid biosynthesis, thereby maintaining a balance between Hcy and Met and producing a molecule that reflects their ratio. It is essential to consider that fluctuations in other central metabolism cycles will also affect this balance. For example, cobalamin deficiencies have been associated with Hcy accumulation in different organisms due to alterations in the folate cycle (42). When the latter is altered, tRNA_i_ formylation is reduced, affecting Met use in translation and contributing to an unbalanced Hcy/Met ratio (43). The Hcy/Met balance can also be affected by the methionine cycle, where Met is used as a substrate in the synthesis of SAM, and SAH recycling produces more Hcy (Figure 6). These observations suggest that HTL plays the role of a signal molecule whose concentration is a consequence of the state of different areas in central metabolism and which needs to reach a threshold concentration to promote the transition from log to exponential growth.

### Homocysteine thiolactone signaling via SdiA

The core of the AI-I-based sensing mechanism consists of a LuxI-type synthase and a LuxR-type receptor, which catalyze the acylation and lactonization reaction between SAM and an acyl carrier protein (ACP), producing an AHL and recognizing and triggering a response to a threshold concentration of ligand, respectively (44). However, *E. coli* and a large group of bacteria lacking AHL synthase have orphan LuxR-type receptors.

The high conservation of residues in the ligand and DNA binding domains and their role in regulating several phenotypes show that they do not correspond to pseudogenes; these receptors recognize AHLs and other signals produced by different bacteria for interspecies communication (45, 46). Our genetic and biochemical analysis indicates that the presence of SdiA is also necessary to respond to HTL in the regulation of lag time (Figure 3B) and that, as a consequence of their interaction (Figure 4), SdiA exerts its role as a transcriptional factor favoring the expression of genes such as *gadY* and *rmf* (Figure 3E). These data do not contradict that *E. coli* uses SdiA to eavesdrop on its neighbors, rather our data show that SdiA can also sense endogenous HTL produced by *E. coli* to regulate the transition from the lag phase to the exponential phase of growth. This phenotype was not observed for other ligands, such as C8-HSL and 3OC8-HSL (Figure 5A).

Due to the potential use of AHLs for bacterial control in both healthcare and biotechnological contexts, some studies have evaluated libraries of AHL homologs for their ability to activate both QS receptors with a known ligand and orphan receptors. These works have revealed that LuxR-type receptors are capable of recognizing molecules where the l lactone ring is replaced by a thiolactone ring, suggesting that the interaction between residues W95 and F100 (Figure 3D), conserved in SdiA and other LuxR homologs, is conserved with the sulfur of thiolactones (47, 48). Structural studies have shown that SdiA has selectivity for short-chain AHLs because the ligand binding pocket cannot accommodate aliphatic chains longer than eight carbons, a requirement met by HTL (49). Since intracellular HTL originates from the L-Hcy editing reaction, all HTL produced by MetRS will correspond to the L-HTL enantiomer, which our data suggests is stereospecific for SdiA activation. This specificity allows us to reinterpret some of the experiments in which, in the search for ligands for LuxR-like receptors from *E. coli* and *S. enterica*, the activation of SdiA was analyzed using DL-HTL, for which no effect was observed (50). Other molecules unrelated to the general structure of AHLs have been investigated as ligands of LuxR homologs, such as 1-octanoyl-rac-glycerol (monocaprylin). This molecule has been proposed as an endogenous ligand for SdiA because they were copurified, with monocaprylin occupying the ligand binding pocket (51). Because this molecule lacks a lactonic ring structure, part of the previously described interactions between SdiA and C8-HSL disappear. The authors proposed that these missing interactions are compensated for by glycerol molecules that fill the gaps in the binding pocket. On the other hand, it has been shown that this molecule interacts with the membrane, and its accumulation has antimicrobial properties due to the destabilization of the membrane (52). Considering that no biological effect on SdiA has been observed in the presence of 1-octanoyl-rac-glycerol, it is not clear that it is a functional ligand of SdiA.

### Homocysteine thiolactone provides a general signal for cell signaling

Since Hcy and MetRS are found in all bacteria, the accumulation of HTL in response to changes in amino acid homeostasis is expected to be a common occurrence. This allows us to suggest that a similar mechanism to that observed in *E. coli* could occur in bacteria without AI-I synthetases, where virulence has been associated with SdiA activity, such as *Salmonella spp*., *Klebsiella spp*., *Shigella spp*., among others (9, 10). An exciting question arises when we integrate this new element into bacteria that actively produce autoinducers for QS. Recently, HTL was proposed as an anti-QS molecule because it inhibits QS in the bioreporter *Chromobacterium violaceum* and *P. aeruginosa* (37). Although it is not clear whether the observed anti-QS effects were due to the use of the racemic mixture DL-HTL, *P. aeruginosa* has the receptors LasR and RhlR that are part of the LuxR protein family, where HTL could compete with the species-specific ligands produced by the bacteria. Our findings link a ubiquitous mechanism for MetRS-catalyzed HTL synthesis to bacteria previously considered silent due to the lack of homologs of the LuxI synthase, which gives them voice. This resolves important questions about bacterial communication and fosters new ideas about the interaction of HTL with the canonical QS elements.

## Materials and Methods

### Strains and culture media

All experiments in this work were performed using wild-type (WT) or mutants of the parental strain *E. coli* K-12 BW30270. Strains and plasmids are listed in Table S1. Strains were cultured in lysogeny broth (LB) media (1% tryptone, 0.5% yeast extract, and 0.5% NaCl), M9 media (47.7 mM Na_2_HPO_4_, 22.0 mM KH_2_PO_4_, 8.6 mM NaCl, 18.7 mM NH_4_Cl, 2 mM MgSO_4_, 0.1 mM CaCl_2_, thiamine 1μg/mL, FeSO_4_*7H_2_O 0.5μg/mL, and 0.4% Glucose). When indicated, amino acids (40μg/mL each, except Met), Met (320μM), Hcy (320μM), HTL (320μM or 500μM), N-octanoyl-L-homoserine lactone (C8-HSL; 320μM), N-(3-Oxooctanoyl)-L-homoserine lactone (3OC8-HSL; 320μM), DL-HTL (320μM), ampicillin (100 μg/ml), arabinose 0.4%, or glucose 0.4%, were added to the culture media.

### Growth curves

*E. coli* overnight (ON) cultures in LB media were washed 3 times with fresh M9 media and diluted to an OD600nm ∼0.1 in M9. Aliquots of 10 μL were transferred to a 96-well microplate with 190 μL of M9 in each well. Growth was monitored on a SPARK microplate reader, Tecan, by measuring absorbance at 600 nm, at 37°C, with constant shaking, alternating between orbital shaking (142 rpm, 6 mm width) and linear shaking (296 rpm, 6 mm width). Doubling and lag times were calculated from the slope and x-intercept of the exponential growth curve, respectively.

### HTL quantification

*E. coli* cultures were performed following the same protocol as the growth curves, but in a volume of 12 mL of M9. At each point of the curve, 1 mL of culture was taken, which was used to measure the OD600, then centrifuged for 2 min at 16000g (room temperature), and 800 μL of its supernatant was stored at -80°C until analysis. Aliquots of 100 μL were mixed with 5 μL of 4 M NaOH and incubated for 15 min at 42°C, to hydrolyze thiolactone into homocysteine. This solution was cooled to room temperature and then neutralized with 5 μL of 4 M HCl. 100 μL of this solution was transferred to a black 96-well microplate containing 100 μL of 4 mM OPA in PBS, and immediately transferred to a plate reader to measure the fluorescence intensity (Ex. 370 ± 20 nm, Em. 420 ± 20 nm) over a period of 30 min every 20 s (21, 23). Triplicates of HTL hydrolysis and OPA reaction were performed for each sample.

### Cloning and Mutation Protocols

PON1 is a mammalian serum paraoxonase whose expression in *E. coli* has been limited due to its aggregation and lack of glycosylations. In this study, we used the G3C9 variant, a rabbit PON1 developed to improve its expression in *E. coli* (24). PON1-G3C9 was amplified from plasmid pG3C9 using primers EcoRI-PON1 and KpnI-PON1, and cloned into the low-copy, arabinose-inducible plasmid pBAD30 using restriction enzymes EcoRI and KpnI. sdiA was amplified from genomic DNA of *E. coli* BW30270 using primers EcoRI-SdiA-fw and HindIII-SdiA-H8-rv (Table S2), which added an 8HisTag at the N-terminus (pBAD-SdiA-H8), and cloned into plasmid pBAD30. *metG*Δ*sdiA* was constructed by the Red-swap method (53) using primers sdiA(H1+P1) and sdiA(H2+P2) (Supplementary Table S2) and plasmid pKD4 as template for amplification of a KanR resistance cassette flanked by FRT sites. BW30270 was transformed with pKD46 to catalyze recombination. All constructs were verified by Sanger sequencing.

### RNA extractions

E. coli ON cultures in LB medium were washed 3 times with fresh M9 medium and incubated for 1 h at 37°C with shaking to adjust to the media change. The cultures were split, and half was supplemented with 320 μM HTL. 1 ml of each bacterial culture was pelleted for 1 min at 12,000 g. The pellet was resuspended in 50 μl of lysis buffer (83 mM Tris HCl, pH 6.8, 18 mM EDTA pH 8, 1.7% SDS, and 1.6% 2-mercaptoethanol) and incubated for 3 min at 37°C. 1.5 ml of TRIzol was added, and total RNA was extracted following the manufacturer recommended protocol.

### qRT-PCRs

Total RNA (5 μg) was treated with TURBO™ Dnase (Invitrogen) followed by a retrotranscription reaction with 2.4 μg RNA, random primers, and SuperScript IV (Invitrogen) according to the manufacturer protocol. The resulting cDNA was used for real-time PCR, with primers specific for *gadY* or *rmf* genes (Table S2) and SsoAdvanced Universal SYBR Green Supermix (Bio-Rad), in a CFX Connect Real-Time PCR Detection System (Bio-Rad). Changes in expression level in M9 or M9+HTL media were represented as Log_2_ Fold Change (ΔCt_M9_-ΔCt_HTL_) (54).

### SdiA purification

SdiA purification was performed using *E. coli* BL21-DE3/XJ1 (Table S1). One-liter LB cultures supplemented with Amp were inoculated with 5% ON preinoculum and grown for 3 h at 37°C. SdiA-H8 was induced with 10 mM Ara for 3 h under the same growth conditions. Cells were harvested by centrifugation (4000g, 4°C) and pellets were stored at -80°C until use. Bacteria were resuspended in 10 mL of lysis buffer (25 mM sodium phosphate buffer, pH 7.4, 150 mM NaCl, 1% Triton-X100, and protease inhibitor cocktail) and sonicated on ice. The lysate was centrifuged (25,000 g, 30 min, 4°C) and the recovered supernatant was fractionated by protein precipitation between 1.2 and 1.6 M ammonium sulfate. The pellet was resuspended in 10 ml of lysis buffer, filtered (0.45 μm, nitrocellulose), and injected into ÄKTA start FPLC (Cytiva). Due to the charge of SdiA-H8, a cation exchange column (1 mL, HiTrap Q FF, Cytiva) and a nickel column (1 mL, HisTrap FF, Cytiva) were connected in tandem for protein loading. After disconnection of the exchange column, the nickel column was washed with buffer A (25 mM sodium phosphate, pH 7.4, 25 mM imidazole, and 800 mM NaCl), buffer B (25 mM sodium phosphate, pH 7.4, 115 mM imidazole, and 150 mM NaCl), and eluted with buffer E (25 mM sodium phosphate, pH 7.4, 500 mM imidazole, and 150 mM NaCl). SdiA-H8 was dialyzed in 10,000 MWCO dialysis tubing (Slide-A-Lyser®Mini dialysis units, Thermo Scientific) in HEPES Buffer (20mM HEPES, pH 7.4, 100mM NaCl). The purity of SdiA-H8 was analyzed by SDS-PAGE and Western blot anti-Hisx6.

### Differential Scanning Fluorimetry

Purified SdiA-H8 (final concentration 3.1 μM) and ligand (final concentration 5 mM) were mixed and incubated for 30 min at 4°C. After the addition of SYPRO Orange fluorophore (Sigma), the reaction was immediately transferred to a precooled CFX Opus 96 real-time PCR system. Cycles of 20 s incubation and fluorescence measurement (FRET scan mode) were performed in a thermal scan from 4 to 90°C (+0.5°C/cycle). The apparent melting temperature (Tm_a_) and maximum fluorescence (Max) were determined from the derivatives of the fluorescence curves, using a protein:fluorophore molar ratio of 1:2.6, while the fluorophore quenching was determined with a ratio of 1:2.26. The same procedure was applied for Hcy, Met, and ligand vehicle (H_2_O) controls.

## Supplementary Material

### Translation Efficiency Analyses

*E. coli* strains were transformed with the plasmid pMet-*gfp*, which contains an arabinose-inducible operon composed of a superfold green fluorescent protein (sfGFP) variant with 4 Met repeats at the sixth codon, followed by a modified mCherry fluorescent protein (55). Bacteria from an ON culture were inoculated into an M9 media supplemented with amino acids and ampicillin, and grown at 37°C with constant shaking. When bacteria reached the mid-log phase (OD600 ∼0.4–0.6), a 50 μl aliquot was transferred to a 96-well optical bottom plate containing 150 μl of fresh M9 medium supplemented with arabinose (final concentration 0.4%) and Met where indicated. The plates were incubated at 37°C with constant shaking in a microplate reader (SPARK, Tecan), where absorbance (600 nm), fluorescence intensity of green fluorescent protein (GFP; Ex. 480 ± 4.5 nm, Em. 515 ± 10 nm) and mCherry (Ex. 555 ± 4.5 nm, Em. 600 ± 10 nm) were periodically recorded.

**Figure S1.**
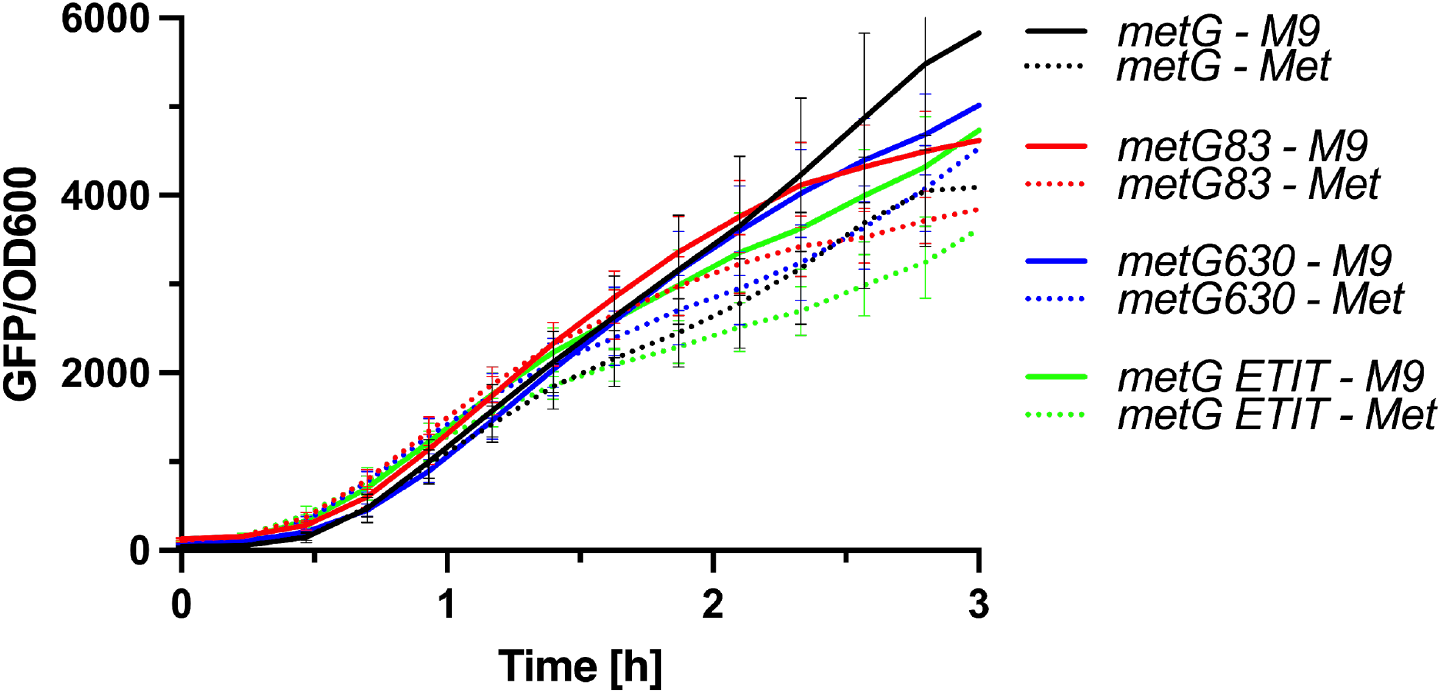
Comparison of protein synthesis between *metG* mutants. Strains *metG* (wild-type), *metG83, metG630*, and *metG ETIT* were transformed with a reporter system encoding the GFP superfolder (GFP) downstream of the inducible PBAD promoter. Kinetics of fluorescence intensity normalized by OD600 (GFP/OD600) was performed on M9, where glucose is replaced by glycerol (M9) and medium supplemented with Met (dotted lines).

**Table S1.** Escherichia coli strains and plasmids used in this study.

| Strain or plasmid | Relevant Features | Source or Reference |
| --- | --- | --- |
| *Strains* |  |  |
| *metG* | BW30270; MG1655 rph+ (wild-type strain) | Coli Genetic Stock Center |
| *metG 630* | BW30270; *metG* mutant (-4TGAT2183) | Wood et al., 2024 |
| *metG 83* | BW30270; *metG* mutant (G263S, S264F) | Wood et al., 2024 |
| *metG ETIT* | BW30270; *metG* mutant (ΔGAAACCATCACC2011) | Wood et al., 2024 |
| *metG*Δ*sdiA* | BW30270 Δ*sdiA*::*KanR* | This study |
| *Plasmids* |  |  |
| pG3C9 | pET32b(+)-PON1<sub>G3C9</sub> | Aharoni et al., 2004 |
| pPON1 | pBAD30-PON1<sub>G3C9</sub> | This study |
| pKD4 | Ori<sub>R6K</sub>, KanR | Datsenko and Wanner, 2000 |
| pKD46 | oriR101 w/repA101ts, AmpR | Datsenko and Wanner, 2000 |
| pMet-*gfp* | PBAD30SFIT-sfgfp(4xMet), mCherry | Leiva et al., 2022 |

**Table S2.**
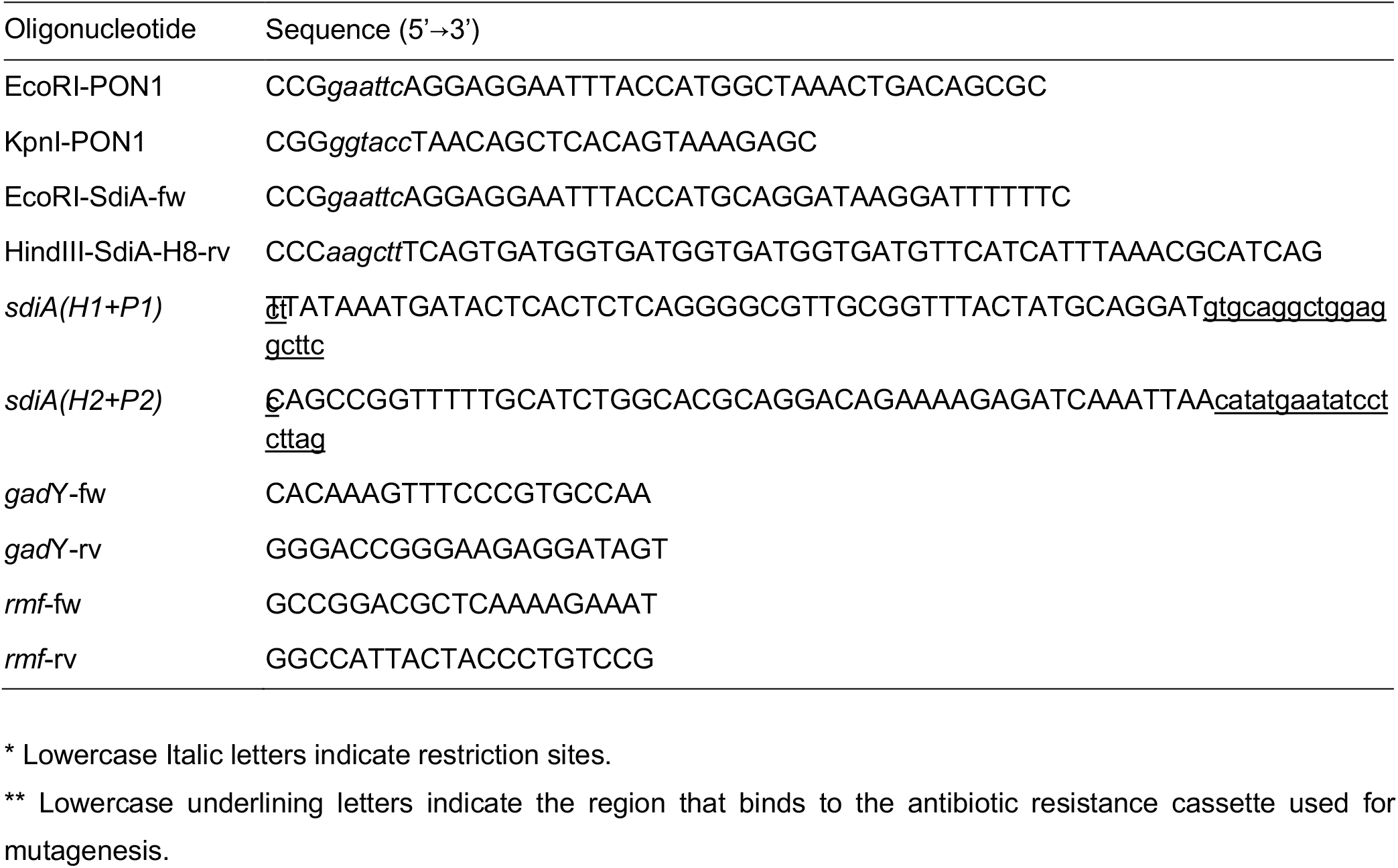
Primers used in this study.

## Acknowledgments

The authors wish to extend their thanks to Dr. Nicholas Backes for help with strain construction, and to reviewers of a previous version of this manuscript for their helpful and constructive comments.

## References

1. S. Mukherjee, B. L. Bassler, Bacterial quorum sensing in complex and dynamically changing environments. Nat. Rev. Microbiol. 17, 371–382 (2019).

2. K. Papenfort, B. L. Bassler, Quorum sensing signal–response systems in Gram-negative bacteria. Nat. Rev. Microbiol. 14, 576–588 (2016).

3. W. R. J. D. Galloway, J. T. Hodgkinson, S. D. Bowden, M. Welch, D. R. Spring, Quorum Sensing in Gram-Negative Bacteria: Small-Molecule Modulation of AHL and AI-2 Quorum Sensing Pathways. Chem. Rev. 111, 28–67 (2011).

4. M. J. Bottomley, E. Muraglia, R. Bazzo, A. Carfì, Molecular insights into quorum sensing in the human pathogen Pseudomonas aeruginosa from the structure of the virulence regulator LasR bound to its autoinducer. J. Biol. Chem. 282, 13592–13600 (2007).

5. Y. Yao, et al., Structure of the Escherichia coli Quorum Sensing Protein SdiA: Activation of the Folding Switch by Acyl Homoserine Lactones. J. Mol. Biol. 355, 262–273 (2006).

6. C. Bez, A.M. Geller, A. Levy, V. Venturi, Cell-Cell Signaling Proteobacterial LuxR Solos: a Treasure Trove of Subgroups Having Different Origins, Ligands, and Ecological Roles. mSystems 8:e01039–22 (2023).

7. J.-H. Lee, Y. Lequette, E. P. Greenberg, Activity of purified QscR, a Pseudomonas aeruginosa orphan quorum-sensing transcription factor. Mol. Microbiol. 59, 602–609 (2006).

7a. Y. Lequette Y et al. A distinct QscR regulon in the Pseudomonas aeruginosa quorum-sensing circuit. J Bacteriol. 188, 3365–3370 (2006).

7b. A.O. Brachmann et al., Pyrones as bacterial signaling molecules. Nat. Chem. Biol., 9, 573–578 (2013).

7c. S. Brameyer et al., Dialkylresorcinols as bacterial signaling molecules. Proc. Natl. Acad. Sci. USA, 112, 572–577 (2015).

8. X. Ma, et al., SdiA Improves the Acid Tolerance of E. coli by Regulating GadW and GadY Expression. Front. Microbiol. 11, 1078 (2020).

9. T. Pacheco, et al., SdiA, a Quorum-Sensing Regulator, Suppresses Fimbriae Expression, Biofilm Formation, and Quorum-Sensing Signaling Molecules Production in Klebsiella pneumoniae. Front. Microbiol. 12, 597735 (2021).

10. J. L. Smith, P. M. Fratamico, X. Yan, Eavesdropping by Bacteria: The Role of SdiA in Escherichia coli and Salmonella enterica Serovar Typhimurium Quorum Sensing. Foodborne Pathog. Dis. 8, 169–178 (2011).

10a. B.M.M. Ahmer, Cell-to-cell signalling in Escherichia coli and Salmonella enterica. Mol. Micro. 52, 933–945 (2004).

10b. A. Schwieters & B.M.M. Ahmer, Identification of new SdiA regulon members of Escherichia coli, Enterobacter cloacae, and Salmonella entericaserovars Typhimurium and Typhi. Microbiol. Spectr., e01929–24 (2024).

11. M. A. Rubio Gomez, M. Ibba, Aminoacyl-tRNA synthetases. RNA N. Y. N 26, 910–936 (2020).

12. M. P. Ferla, W. M. Patrick, Bacterial methionine biosynthesis. Microbiology 160, 1571–1584 (2014).

13. H. Jakubowski, Aminoacyl-tRNA synthetases and the evolution of coded peptide synthesis: the Thioester World. FEBS Lett. 590, 469–481 (2016).

14. H. Jakubowski, Metabolism of Homocysteine Thiolactone in Human Cell Cultures. J. Biol. Chem. 272, 1935–1942 (1997).

15. H. Jakubowski, The molecular basis of homocysteine thiolactone-mediated vascular disease. Clin. Chem. Lab. Med. 45 (2007).

16. A. S. Chubarov, Homocysteine Thiolactone: Biology and Chemistry. Encyclopedia 1, 445–459 (2021).

17. W. N. Wood, M. A. Rubio, L. E. Leiva, G. J. Phillips, M. Ibba, Methionyl-tRNA synthetase synthetic and proofreading activities are determinants of antibiotic persistence. Front. Microbiol. 15, 1384552 (2024).

18. G. Sezonov, D. Joseleau-Petit, R. D’Ari, Escherichia coli physiology in Luria-Bertani broth. J. Bacteriol. 189, 8746–8749 (2007).

19. H. Jakubowski, R. Głowacki, “Chemical Biology of Homocysteine Thiolactone and Related Metabolites” in Advances in Clinical Chemistry, (Elsevier, 2011), pp. 81–103.

20. M. Sikora, H. Jakubowski, Homocysteine editing and growth inhibition in Escherichia coli. Microbiology 155, 1858–1865 (2009).

21. H. Jakubowski, The determination of homocysteine-thiolactone in biological samples. Anal. Biochem. 308, 112–119 (2002).

22. H. Jakubowski, Proofreading in vivo: editing of homocysteine by methionyl-tRNA synthetase in Escherichia coli. Proc. Natl. Acad. Sci. 87, 4504–4508 (1990).

23. Y. Mukai, T. Togawa, T. Suzuki, K. Ohata, S. Tanabe, Determination of homocysteine thiolactone and homocysteine in cell cultures using high-performance liquid chromatography with fluorescence detection. J. Chromatogr. B 767, 263–268 (2002).

24. A. Aharoni, et al., Directed evolution of mammalian paraoxonases PON1 and PON3 for bacterial expression and catalytic specialization. Proc. Natl. Acad. Sci. 101, 482–487 (2004).

25. M. Sarkar, et al., Solubilization and Humanization of Paraoxonase-1. J. Lipids 2012, 1–13 (2012).

26. R. F. Anderson, J. E. Packer, The radiolysis of aqueous solutions of homocysteinethiolactone hydrochloride. Int. J. Radiat. Phys. Chem. 6, 33–46 (1974).

27. H. Jakubowski, Mechanism of the Condensation of Homocysteine Thiolactone with Aldehydes. Chem. – Eur. J. 12, 8039–8043 (2006).

28. S. Youssef-Saliba, A. Milet, Y. Vallée, Did Homocysteine Take Part in the Start of the Synthesis of Peptides on the Early Earth? Biomolecules 12, 555 (2022).

29. V. Du Vigneaud, W. I. Patterson, M. Hunt, OPENING OF THE RING OF THE THIOLACTONE OF HOMOCYSTEINE. J. Biol. Chem. 126, 217–231 (1938).

30. K. E. Kram, S. E. Finkel, Rich Medium Composition Affects Escherichia coli Survival, Glycation, and Mutation Frequency during Long-Term Batch Culture. Appl. Environ. Microbiol. 81, 4442–4450 (2015).

31. R. Sánchez-Clemente, et al., Study of pH Changes in Media during Bacterial Growth of Several Environmental Strains in Environment, Green Technology, and Engineering International Conference, (MDPI, 2018), p. 1297.

32. H. Yoshida, T. Shimada, A. Ishihama, Coordinated Hibernation of Transcriptional and Translational Apparatus during Growth Transition of Escherichia coli to Stationary Phase. mSystems 3, e00057–18 (2018).

33. F. H. Niesen, H. Berglund, M. Vedadi, The use of differential scanning fluorimetry to detect ligand interactions that promote protein stability. Nat. Protoc. 2, 2212–2221 (2007).

34. T. Wu, et al., Protocol for performing and optimizing differential scanning fluorimetry experiments. STAR Protoc. 4, 102688 (2023).

35. J. Alesio, G. D. Bothun, Differential scanning fluorimetry to assess PFAS binding to bovine serum albumin protein. Sci. Rep. 14, 6501 (2024).

35a. B. Michael, J.N. Smith, S. Swift, F. Heffron, B.M.M. Ahmer, SdiA of Salmonella enterica Is a LuxR Homolog That Detects Mixed Microbial Communities. J Bacteriol. 183, 5733–5742 (2001).

35b. J.N. Smith & B.M.M. Ahmer, Detection of other microbial species by Salmonella: expression of the SdiA regulon. J.Bacteriol. 185, 1357–1366 (2003).

36. A. R. McCready, J. E. Paczkowski, B. R. Henke, B. L. Bassler, Structural determinants driving homoserine lactone ligand selection in the Pseudomonas aeruginosa LasR quorum-sensing receptor. Proc. Natl. Acad. Sci. 116, 245–254 (2019).

37. Z.-P. Ma, et al., Anti-quorum Sensing Activities of Selected Coral Symbiotic Bacterial Extracts From the South China Sea. Front. Cell. Infect. Microbiol. 8, 144 (2018).

38. H. Jakubowski, Homocysteine Modification in Protein Structure/Function and Human Disease. Physiol. Rev. 99, 555–604 (2019).

39. O. Fridman, A. Goldberg, I. Ronin, N. Shoresh, N. Q. Balaban, Optimization of lag time underlies antibiotic tolerance in evolved bacterial populations. Nature 513, 418–421 (2014).

40. E. Şimşek, M. Kim, Power-law tail in lag time distribution underlies bacterial persistence. Proc. Natl. Acad. Sci. 116, 17635–17640 (2019).

41. C. Vulin, N. Leimer, M. Huemer, M. Ackermann, A. S. Zinkernagel, Prolonged bacterial lag time results in small colony variants that represent a sub-population of persisters. Nat. Commun. 9, 4074 (2018).

42. A. Mahajan, et al., Effect of imbalance in folate and vitamin B12 in maternal/parental diet on global methylation and regulatory miRNAs. Sci. Rep. 9, 17602 (2019).

43. K. Lahry, A. Gopal, S. Sah, R. A. Shah, U. Varshney, Metabolic Flux of N10-Formyltetrahydrofolate Plays a Critical Role in the Fidelity of Translation Initiation in Escherichia coli. J. Mol. Biol. 432, 5473–5488 (2020).

44. W. T. Watson, T. D. Minogue, D. L. Val, S. B. Von Bodman, M. E. A. Churchill, Structural Basis and Specificity of Acyl-Homoserine Lactone Signal Production in Bacterial Quorum Sensing. Mol. Cell 9, 685–694 (2002).

45. A. V. Patankar, J. E. González, Orphan LuxR regulators of quorum sensing. FEMS Microbiol. Rev. 33, 739–756 (2009).

46. G. Xu, Evolution of LuxR solos in bacterial communication: receptors and signals. Biotechnol. Lett. 42, 181–186 (2020).

47. C. E. McInnis, H. E. Blackwell, Thiolactone modulators of quorum sensing revealed through library design and screening. Bioorg. Med. Chem. 19, 4820–4828 (2011).

48. M. J. Styles, et al., Chemical Control of Quorum Sensing in E. coli : Identification of Small Molecule Modulators of SdiA and Mechanistic Characterization of a Covalent Inhibitor. ACS Infect. Dis. 6, 3092–3103 (2020).

49. T. Kim, et al., Structural insights into the molecular mechanism of Escherichia coli SdiA, a quorum-sensing receptor. Acta Crystallogr. D Biol. Crystallogr. 70, 694–707 (2014).

50. J. C. A. Janssens, et al., Synthesis of N -Acyl Homoserine Lactone Analogues Reveals Strong Activators of SdiA, the Salmonella enterica Serovar Typhimurium LuxR Homologue. Appl. Environ. Microbiol. 73, 535–544 (2007).

51. Y. Nguyen, et al., Structural and Mechanistic Roles of Novel Chemical Ligands on the SdiA Quorum-Sensing Transcription Regulator. mBio 6, e02429–14 (2015).

52. M. Hyldgaard, D. S. Sutherland, M. Sundh, T. Mygind, R. L. Meyer, Antimicrobial Mechanism of Monocaprylate. Appl. Environ. Microbiol. 78, 2957–2965 (2012).

53. K. A. Datsenko, B. L. Wanner, One-step inactivation of chromosomal genes in Escherichia coli K-12 using PCR products. Proc. Natl. Acad. Sci. 97, 6640–6645 (2000).

54. K. J. Livak, T. D. Schmittgen, Analysis of Relative Gene Expression Data Using Real-Time Quantitative PCR and the 2−ΔΔCT Method. Methods 25, 402–408 (2001).

55. L. E. Leiva, et al., Oxidative stress strongly restricts the effect of codon choice on the efficiency of protein synthesis in Escherichia coli. Front. Microbiol. 13, 1042675 (2022).

